# Multiplexed microfluidic screening of bacterial chemotaxis

**DOI:** 10.1101/2022.11.30.518228

**Authors:** Michael R. Stehnach, Richard J. Henshaw, Sheri A. Floge, Jeffrey S. Guasto

## Abstract

Microorganism sensing of and responding to ambient chemical gradients regulates a myriad of microbial processes that are fundamental to ecosystem function and human health and disease. The development of efficient, high-throughput screening tools for microbial chemotaxis is essential to disentangling the roles of diverse chemical compounds and concentrations that control cell nutrient uptake, chemorepulsion from toxins, and microbial pathogenesis. Here, we present a novel microfluidic multiplexed chemotaxis device (MCD) which uses serial dilution to simultaneously perform six parallel bacterial chemotaxis assays that span five orders of magnitude in chemostimulant concentration on a single chip. We first validated the dilution and gradient generation performance of the MCD, and then compared the measured chemotactic response of an established bacterial chemotaxis system (*Vibrio alginolyticus*) to a standard microfluidic assay. Next, the MCD’s versatility was assessed by quantifying the chemotactic responses of different bacteria (*Psuedoalteromonas haloplanktis, Escherichia coli*) to different chemoattractants and chemorepellents. The MCD vastly accelerates the chemotactic screening process, which is critical to deciphering the complex sea of chemical stimuli underlying microbial responses.

## I. INTRODUCTION

Motile cells of all types navigate complex environments through the detection of and response to chemical signals via chemotaxis [1–4]. This fundamental survival mechanism regulates countless biological processes, such as microbial foraging in marine environments [5, 6] and reproduction [7]. Consequently, considerable effort has been invested into the study of microbial chemotaxis [8, 9] to better understand their chemotactic motility [10], detection sensitivity [11], and transport [12]. Microfluidic devices have become an indispensable platform for disentangling the intricacies of microbial chemotaxis by virtue of their precise control over the chemical environment at scales relevant to swimming cells [8, 13]. Specifically, microfluidics have been employed to physically model a range of chemical landscapes, such as nutrient patches [6], and provide highly tunable concentration profiles [14, 15]. Microfluidics have been broadly applied across microbial systems for both drug-dose response quantification [15] and infectious disease diagnostics [16]. While microfluidic chemotaxis assays have evolved since their inception [13], the vast landscape of potential chemical compounds, combinations of compounds, and concentration gradient conditions that regulate these important processes necessitates the development of new high-throughput devices.

Faced with a broad range of chemostimulant concentrations and gradients in their natural environment [3], microorganisms, specifically prokaryotes, have evolved exquisite chemosensing abilities with variable degrees of specificity to nutrients, dissolved resources, toxins, and signaling molecules [1–3]. Some bacteria exhibit a dynamic sensing range spanning five orders of magnitude [10, 17, 18] and can detect nano-molar attractant concentrations [11], while marine invertebrate spermatozoa have a reported detection limit approaching the femto-molar scale [19]. Quantifying the strength of chemotactic responses across varying concentration and concentration gradient conditions presents a key challenge to understanding microbial driven processes, extending far beyond their search for optimal metabolic activity conditions [8]. For example, in marine microbial communities, the natural phycosphere surrounding individual cells [20] contains diverse spectrum and concentration of metabolite and organic material [21], which are taken up by chemotaxing microbes [22]. Viral infection of microbes augments this process and is a principle mechanism [21, 23] for transforming live biomass to readily available organic matter. Lysis [24] and exudation [25] by virus infected cells releases a diverse range and concentration of metabolite and organic material [21]. Furthermore, chemotaxis is essential in initiating bacterial infections and pathogenicity for both animals and plants [26]. For example, in gastric infections pathogenic organisms rapidly colonize surfaces via chemotaxis, where a range of attractants from urea to amino acids and metals are presumed to enable localization and colonization on the host epithelium [27]. Identifying the key metabolites and signaling chemicals which drive microbial chemotaxis necessitates new microfluidic tools capable of probing the wide scope and scale of chemotactic behaviors across a myriad of complex systems.

Microfluidic devices are widely accepted as an indispensable platform for targeted chemotaxis assays by enabling the quantification of both single cell and population-scale responses to precisely-controlled chemical gradients [13, 14]. One class of chemotaxis microfluidic devices, termed stop-flow diffusion, relies on flowing a chemostimulant solution and buffer stream side-by-side in a microchannel. Upon halting the flow a slowly-evolving concentration gradient forms via diffusion (Fig. 1a,b) [5, 6, 11, 13]. Other devices generate steady chemical gradients by utilizing porous materials [13] or micro-well assays to mimic diffusing marine hotspots that entice and trap chemotactic microorganisms [20]. While these well-established assays accurately measure chemotactic motility in physically relevant concentration gradients [6], they largely overlook the potential for high-throughput screening afforded by microfluidic devices. Recently, such high-throughput capabilities have been broadly showcased in other fields through the use of parallelized microfluidics for clinical testing of viruses [16], drug responses [15], and cell profiling [30]. The development of an integrated microfluidic design - comprising parallelized chemotaxis assays on a single chip - would enable high-throughput characterization of microbial chemotactic responses. Relative to time-prohibitive conventional assays, rapid chemotaxis phenotyping could facilitate comparative studies and discoveries across different swimming microorganisms, chemostimulants, and concentration gradient conditions.

**FIG. 1.**
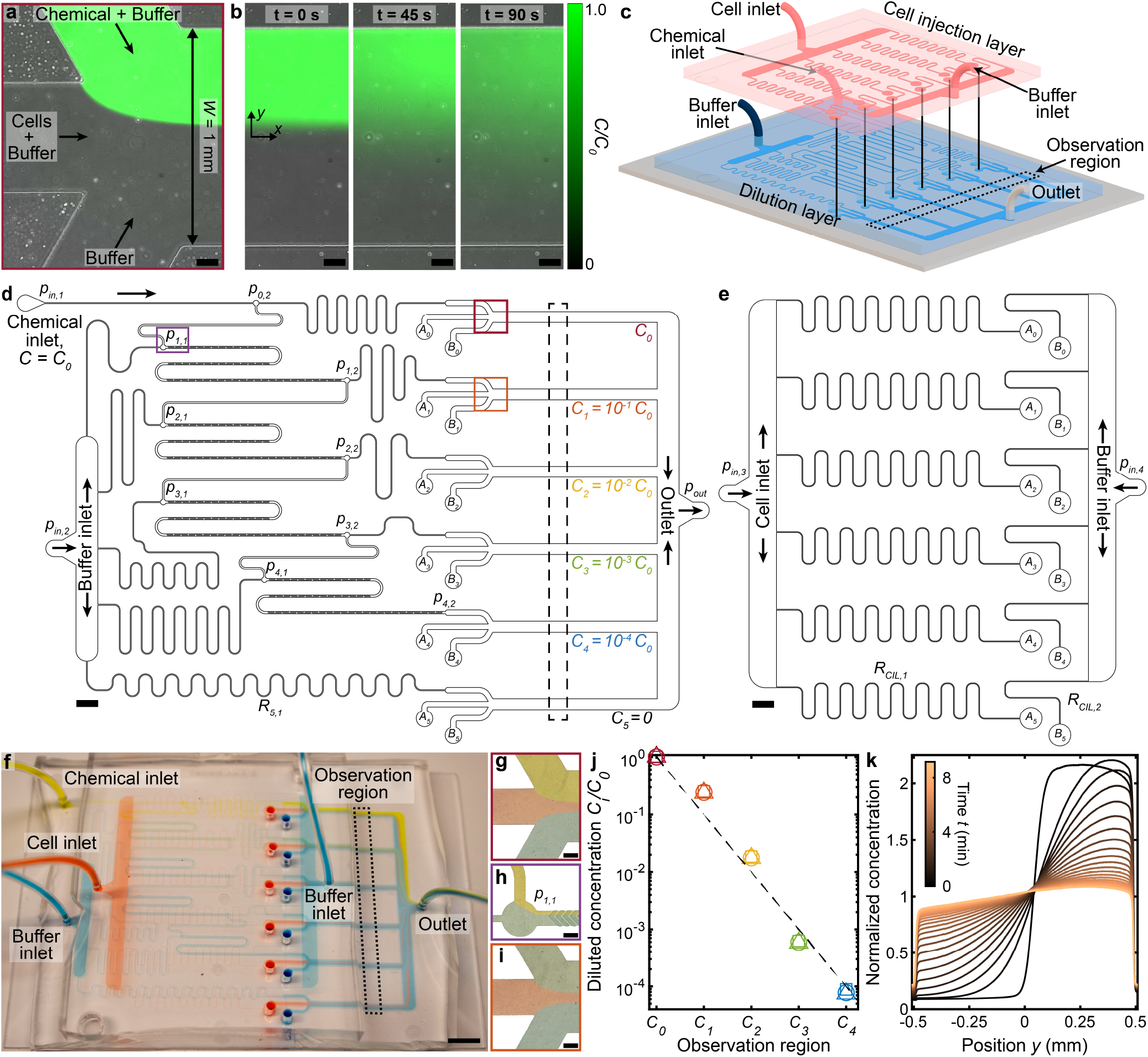
Multiplexed microfluidic device for simultaneous chemotaxis assays. (**a,b**), A microfluidic junction (a) stratifies chemostimulus, cell, and buffer solutions, demonstrated here with fluorescein, DI water, and DI water, respectively [13, 28]. Upon halting the flow (b), diffusion establishes a chemical gradient across the channel, which is repeated at each observation channel in the MCD (d, red and orange boxes). Scale bars, 0.1 mm. (**c**), Assembly of the MCD showing the PDMS dilution layer (blue) and cell injection layer (red) microchannels mounted on a glass slide (grey; **Methods**). (**d**), Scaled drawing of the dilution layer, which receives chemical (pressure, *p*_*in*,1_) and buffer (*p*_*in*,2_) solutions. Initial chemical concentration (*C*_0_) is sequentially diluted 10-fold to each of four additional concentrations (*C*_1-4_), plus a control solution (*C*_5_ = 0). These six chemostimulus solutions are merged separately with additional cell (*A_i_*) and buffer (*B_i_*) solutions from the cell injection layer (e) for chemotaxis assays in respective observation channels (dashed black box, corresponding to c and f). (**e**), Scaled drawing of the cell injection layer which injects a cell suspension (*p*_*in*,3_) and buffer solution (*p*_*in*,4_ = *p*_*in*,3_) into the dilution layer (*A_i_, B_i_*; **Methods**). Scale bars d,e, 2 mm. (**f**), Photograph of the completed MCD with dyed water to visualize the chemical (yellow), cell (red), and buffer (blue) fluid streams in the channel network. Scale bar, 5 mm. (**g**), Stratified chemical (*C*_0_), cell, and buffer solutions in the first observation region (d, red box). (**h**), Dilution of the chemical (*C*_0_) by the buffer prior to mixing in the first micromixer [29] to produce concentration *C*_1_ (d, purple box). (**i**), Stratified chemical solution after initial dilution (*C*_1_, green) in the second observation region (d, orange box). Scale bars g-i, 0.2 mm. (**j**), Measured chemical concentrations (see **Methods**) generated from the dilution microchannels (d) for various driving pressures *p*_*in*,1_ = *p*_*in*,2_ = [100, 150, 200] mbar (square, circle, and triangle, respectively). (**k**), Measured evolution of the chemical gradient (b) produced in the *C*_0_ observation region (Fig. 7; **Methods**) by the MCD shows the chemical diffusion across the channel with increasing time t.

Here, we present a microfluidic multiplexed chemotaxis device (MCD) that enables high-throughput chemotaxis screening of swimming microorganisms to chemical stimuli across concentration gradient conditions that potentially span the microorganism’s entire sensitivity range. The two-layer device architecture comprises a serial dilution layer that produces logarithmically-diluted chemostimulant solutions [15] and a cell injection layer that introduces swimming cells, whilst minimizing both the footprint and operational complexity of the device (Fig. 1c-e). On a single chip, the MCD simultaneously performs six stop-flow diffusion chemotaxis assays (including control), which span five orders of magnitude in chemostimulant concentration. The dilution, mixing, gradient generation, and flow performance are fully characterized (**Methods**), and the MCD is validated against a conventional chemotaxis device for a known marine bacterial chemotaxis system (*Vibrio alginolyticus*). To demonstrate the device’s efficiency, capabilities, and operational flexibility, the MCD is then used to rapidly quantify the chemotactic responses of different microbes to a variety of chemostimulants. Compared to existing microfluidic devices, the MCD enables chemotaxis studies with significantly higher throughput rates, and most importantly facilitates the simultaneous measurement of chemotactic responses across a range of concentration gradient conditions.

## II. RESULTS

### A. Multiplexed microfluidic device as a platform for high throughput chemotaxis screening

To enable rapid and efficient chemotaxis screening of swimming microbes, we designed the multiplexed chemotaxis device (MCD) to perform six chemotaxis assays in parallel on a single microfluidic chip (Fig. 1). The individual assays are based on established stop-flow diffusion methods [6, 13, 28], whereby an initially stratified chemostimulus (concentration, *C_i_*) forms a gradient via diffusion with a buffer (*C* = 0) in an observation channel (Fig. 1a,b), and the response of the swimming cells may be observed and recorded. The MCD (Fig. 1c-i) performs six simultaneous assays comprising five logarithmically decreasing chemical concentrations (*C_i_* = 10^-*i*^*C*_0_, *i* ∈ [0, 4]; Fig. 1j), plus one control (*C*_5_ = 0). The device is fabricated from polydimethylsiloxane (PDMS) in two layers (**Methods**; Fig. 1c-e). The primary function of the dilution layer (Fig. 1d) is to receive two fluid inputs - base chemostimulus solution (*C*_0_) and buffer (*C* = 0) - and passively generate six predefined concentration conditions (*C_i_*) via a serial dilution process [14, 15, 31], which are then dispensed to each of the six observation channels for chemotaxis assays. At each stage of the serial dilution process, the chemostimulus stream is combined with buffer in a 1:9 volumetric flow rate ratio, where efficient mixing of the solutions is necessary for accurate dilution and chemotaxis assays downstream. To ensure sufficiently mixed solutions, herringbone micromixers [29] (Fig. 1d,h and Fig. 5) were incorporated into each dilution stage. These structured microchannel surfaces generate a three-dimensional flow to induce chaotic mixing, and thus significantly reduce the mixing length [29] and the overall footprint of the device (**Methods**; Fig. 5). To achieve the targeted chemostimulus concentrations in the dilution layer, the microfluidic network was designed using hydraulic circuit theory (**Methods**; Fig. 6), and the accuracy of the serial dilution process was experimentally confirmed (Fig. 1j).

The cell injection layer (Fig. 1e) introduces a suspension of swimming microbes and a sheathing buffer solution from two corresponding inlets into the six observation regions of the dilution layer (Fig. 1d, dashed box). In the observation regions, the chemostimulus solution, cell suspension, and buffer streams comprise six standard stop-flow chemotaxis assays (Fig. 1a), where each one incorporates a unique chemostimulus concentration. The observation channel width (W = 1 mm) and height (*H* = 90 *μ*m) are similar to other microfluidic devices [6, 13, 28] and ensure organisms with different sizes can be studied using the MCD. The initially steady flow rates stratify the three fluids in the observation region and localize the cells in a thin band in the channel center with equal width chemostimulus and buffer streams on either side (**Methods**; Fig. 1g,i and Fig. 7). Upon halting the flow, a unique and highly reproducible chemical gradient is formed in each observation channel via diffusion, where the consistency of the transient concentration profiles across all observation channels were confirmed using fluorescence microscopy (**Methods**; Fig. 1k, Fig. 7). The mixing effectiveness, fluid stratification, and gradient formation were validated over a range of applied pressures (approximately 100-200 mbar) and were found to be consistent across all six observation regions (Fig. 5 and Fig. 7). This efficient two-layer architecture reduces the operational complexity of the device by minimizing the total number of fluid inlets (four) and reduces the footprint of the microfluidic chip.

Both layers of the MCD are fabricated from polydimethylsiloxane (PDMS) through standard soft lithography techniques (replica molding; **Methods**). The final microfluidic chip is assembled by plasma bonding the dilution layer to a standard double-wide microscope slide (75 mm × 50 mm × 1mm) and subsequently aligning and bonding the cell injection layer on top (**Methods**; Fig. 1c,f and Fig. 8). This reusable device is driven by a single pressure pump which maintains flow stratification (1-2 minutes) prior to each assay to ensure consistent initial gradient conditions for measuring cell responses. Pump and microscope automation enables the chemostimulus gradient to be reset for rapid replicate measurements. The design and operation of the MCD can accommodate most single-celled microorganisms, and efficient micromixer channels facilitate the use of a wide range of dissolved chemostimulants (Fig. 5) [29]. Further optimization of the device layout could enable the number of observation channels to be expanded, including a broader range or more refined sampling of concentration gradient conditions. Additionally, the serial dilution layer design can be easily modified to produce different concentration scalings (e.g. logarithmic, linear) [15, 31]. The high degree of parallelization for chemotaxis screening, combined with the demonstrated consistency and repeatability of the chemostimulus gradients, represents a significant advance relative to existing microfluidic chemotaxis devices [6, 13, 14].

### B. Validation of MCD performance against conventional chemotaxis assays

Having established the gradient generation performance, the MCD is compared to a conventional chemotaxis assay to (i) validate the chemotaxis measurements against a known microorganism-chemostimulus system, and (ii) demonstrate the high-throughput capability of the MCD in comparison with existing devices. The chemotactic response of the monotrichous marine bacterium *Vibrio alginolyticus* (Fig. 2a) was measured using both the MCD and a single microfluidic gradient generation device (referred to as “single assay”, SA) [28]. *V. alginolyticus* swims with a run-reverse-flick motility pattern, and it was chosen due to its well-documented chemotaxis [32, 34, 35] and prevalence within marine microbial communities [36]. The single assay device has an identical observation channel geometry with three inlets and operates using stop-flow diffusion in the same manner as the MCD, where the chemostimulus, cells, and buffer are initially flow-stratified (Fig. 2b). Upon halting the flow for both assays, the concentration gradient develops via diffusion, and time-lapse microscopy is used to measure the evolution of the spatial cell distribution over time, *t* (Fig. 2c; **Methods**). For example, in a gradient formed by the model chemoattractant serine [32], cells initially confined to a central band migrate toward the attractant, before uniformly sampling the channel (Fig. 2c,d) as the gradient dissipates within *t* ≈ 10 min. Beyond ensuring the uniformity of the gradient generation in the MCD (Fig. 7), control experiments using *V. alginolyticus* with no chemostimulus and with a fixed serine concentration confirmed the consistency of the chemotaxis assays, having no discernable bias across all MCD observation channels (Fig. 2k,l and Fig. 9).

**FIG. 2.**
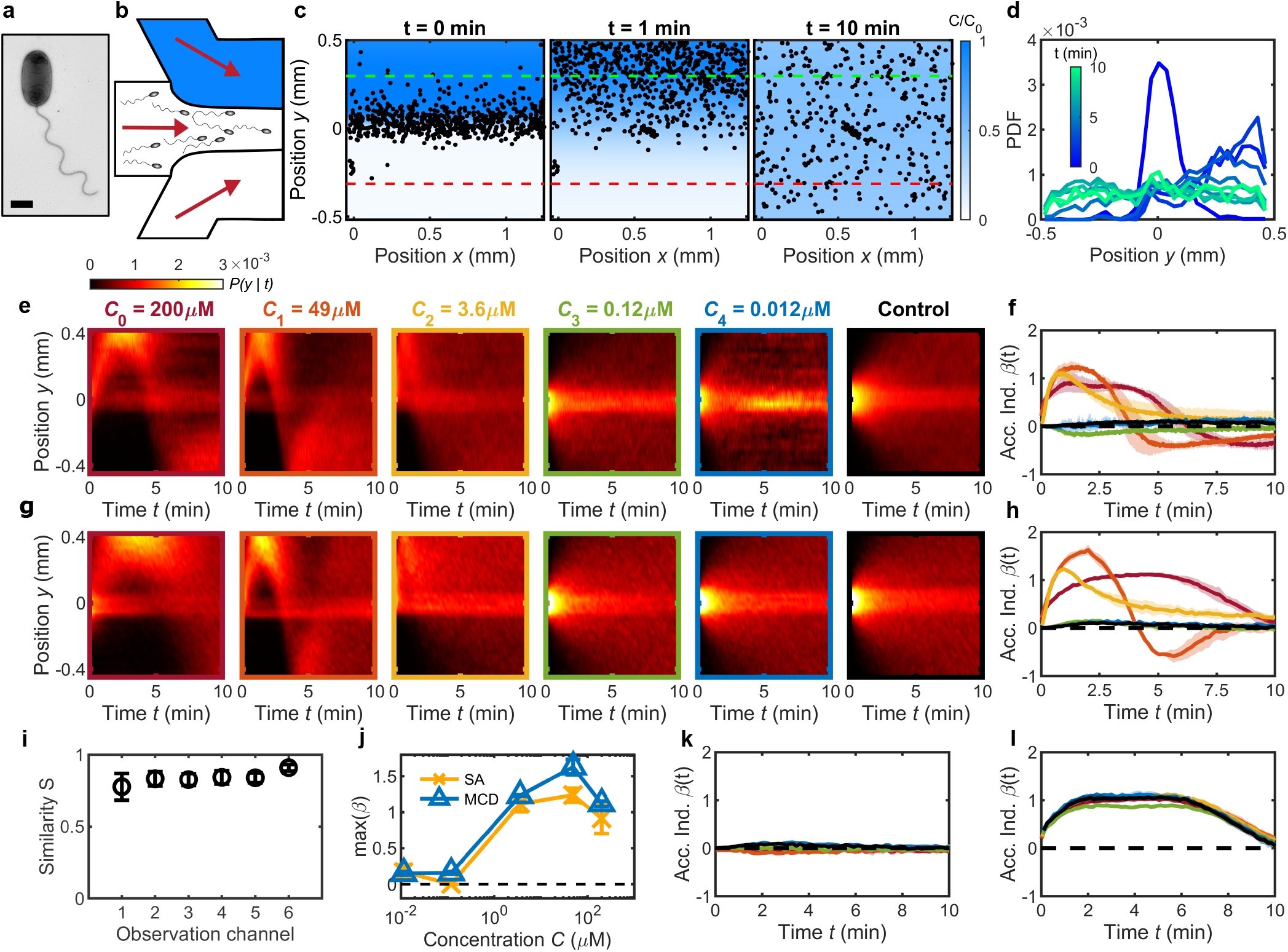
Validation of MCD and measurement of *V. alginolyticus* chemotactic performance toward serine. (**a**), TEM image of *V. alginolyticus* (**Methods**). Scale bar, 1 *μ*m. (**b**), A single chemotaxis assay (SA) with a single conventional microfluidic device flows chemostimulus (top, blue), cell suspension (middle), and buffer (bottom) streams into the observation region (**Methods**). (**c**), SA with chemostimulus (serine, *C* = 100 *μ*M) showing measured cell positions (*V. alginolyticus*, black dots) at various times *t* after initial stratification (*t* = 0 min) relative to the chemostimulus distribution (blue, from measurements in Fig. 1b). Cells migrate up the gradient (t = 1 min) followed by uniform dispersal as the gradient dissipates (*t* = 10 min). Degree of cell accumulation is determined from the number of cells, *N_p,n_*, in a 200 *μ*m wide region on the chemostimulus side (positive; green dashed line) and buffer side (negative; red dashed line), respectively [5, 6]. (**d**), The measured cell distribution across the microchannel evolves over time (from c) and is represented as a conditional probability density of cell position, *P*(*y*|*t*) (shown as a kymograph). (**e**), *P*(*y*|*t*) for *V. alginolyticus* chemotactic response to serine from a series of SA devices having the same geometry as the MCD observation regions (Fig. 1). SA measurements illustrate the transition from positive chemotactic response at high attractant concentration (*C*_0-2_) to no response at low concentration (*C*_3-4_) compared to control (*C*_5_ = 0 *μ*M) [32]. (**f**), Accumulation index, *β*(*t*), for SA measurements from e. g, *P*(*y*|*t*) measured by the MCD under the same conditions of the SA. (**h**), *β*(*t*) measured from g accurately captures the behavior of *V. alginolyticus* to serine compared to SA results (f). (**i**), Sørensen similarity metric [33] comparing e and g, which is calculated at each time point and averaged. (**j**), Comparing MCD and SA peak chemotactic response quantified by max(*β*(*t*)) from f,h. k,l, *β*(*t*) in the absence of a chemical gradient (k; Fig. 9) and for fixed gradients of *C_i_* = 200 *μ*M (l; Fig. 9b) across each observation channel in the MCD indicates no significant bias. Shading in f,h,k,l indicates one standard deviation (N=3). Error bars in i,j are one standard deviation.

From one parallelized assay, the MCD precisely reproduces *V. alginolyticus* chemotactic responses toward various serine concentrations, compared to multiple, conventional single assays (Fig. 2e-h). Chemotactic behavior of the bacteria was measured across a range of manually adjusted attractant concentrations for the single assay, decreasing from *C*_0_ = 200 *μ*M and matching the serial dilution concentrations generated by the MCD (Fig. 1j). For each concentration gradient, the spatial distribution (conditional probability) of cells, *P*(*y*|*t*), across the observation channel, *y*, is shown over time as a kymograph (Fig. 2e,g). For ease of comparison, the aggregate chemotactic response is distilled through the accumulation (or chemotactic) index, *β*(*t*), which quantifies the relative fraction of cells responding to the chemostimulus [5, 6]. The accumulation index is defined as [5, 6]: *β*(*t*) = (*N_p_*(*t*) – *N_n_*(*t*))/(*N_p_*(*T*) + *N_n_*(*T*)), where *N_p,n_*(*t*) are the instantaneous number of cells at time t within predefined positive and negative accumulation regions (Fig. 2c, green and red dashed lines, respectively), and normalized by the total number of cells in these two regions at the final time (*T* ≈ 10 min). The measured cell distributions from the MCD (Fig. 2g) were compared to the single assay results (Fig. 2e) by calculating the Sørensen similarity metric [33] for each attractant concentration, showing excellent statistical agreement between the two assays (Fig. 2i). A comparison of the strength of cell accumulation (Fig. 2j) further emphasizes the strong concordance between the two devices. In particular, *V. alginolyticus* exhibits strong accumulation for high serine concentrations (*C*_0-2_), above the previously reported chemotactic sensitivity threshold of 0.2 *μ*M [32]. Consequently, no discernable response is observed for lower concentrations (*C*_3-4_), which are comparable to *β*(*t*) for the control (*C* = 0). At the highest concentrations (*C*_0-1_), the chemotactic motility exhibits a slight reversal at later times, which is likely due to modification of the concentration gradient via bacterial serine consumption. The persistent central band of cells is due to non-motile bacteria (Fig. 2e,g, near *y* = 0); because these bacteria do not impact the calculation of *β*(*t*), the central band is omitted from future kymographs for clarity. Finally, additional assays with no chemostimulus and with a fixed chemostimulus concentration confirmed the consistency of the chemotaxis assays across all observation regions (Fig. 2k,l and Fig. 9), which is expected from the verified gradient generation performance in each channel (Fig. 7).

These validation assays serve to highlight the dramatically improved efficiency in chemical screening when compared to a standard single assay. In the single assay case, each chemical concentration requires: (i) manually diluting stock solutions, (ii) exchanging peripheral reservoirs for chemicals, and (iii) a new cell suspension for each concentration assay, all of which become extremely costly and time prohibitive, when considering the scope and scale of multi-chemical, concentration, and/or organism panel experiments. The single assay results (Fig. 2e,f) required six different dilutions, cell solutions, and devices, and with three replicates per condition, required 18 individual assays. In contrast, the MCD collected the same data (Fig. 2g,h) in only three automated assays, and did not require culture changes eliminating inconsistencies due to variations in growth media dilution errors or growth conditions. If replacing the cell suspension is necessary, the MCD can be easily reset by simply exchanging the cell suspension with a fresh suspension and restarting the flow. Because it uses a robust serial dilution process and requires the preparation of a single chemostimulus (*C*_0_), the MCD ensures consistent chemical concentrations across different experiments, a crucial factor when working with microorganisms having femto- to nanomolar chemotactic sensitivities [11, 19, 32]. The MCD fully screens both microbe and stimulus pairings with three replicates in ≈ 1 hour with a single cell culture, including the bench time associated with preparing the solutions (Fig. 2h). In contrast, the panel of single assays (Fig. 2f), requires nearly a full working day (≈ 6 – 7 hours). Thus, the MCD is a powerful and much needed tool for large scale chemotactic panel studies, where consistency in stimulus concentration, elimination of biological variability, and the need for efficiency, are essential.

### C. Multiplexed microfluidic device supports high-throughput chemotaxis screening for novel stimuli and various microorganisms

Beyond validation with the single assay device, we demonstrate the efficacy of the MCD by examining the response of *V. alginolyticus* to both a known repellent and novel chemostimulus. Chemorepellents serve as an early warning sign for microorganisms to evade predators and toxins for survival [39]. A single MCD assay reveals that *V. alginolyticus* exhibits negative chemotaxis (*β* < 0) to the model repellent phenol [37] with an observed detection threshold on the order of *C* = 1 – 10 *μ*M (Fig. 3a,b). Separately, the amino acid leucine has been identified as an important metabolite in human health [40] and marine environments [41, 42], and it serves as a measure of prokaryote heterotrophic activity from viral lysis in deep-sea environments [43]. Previously reported as an attractant for marine prokaryotes [44], we verify the positive chemotactic response of *V. alginolyticus* to leucine through rapid chemotaxis screening using the MCD (Fig. 3c,d).

**FIG. 3.**
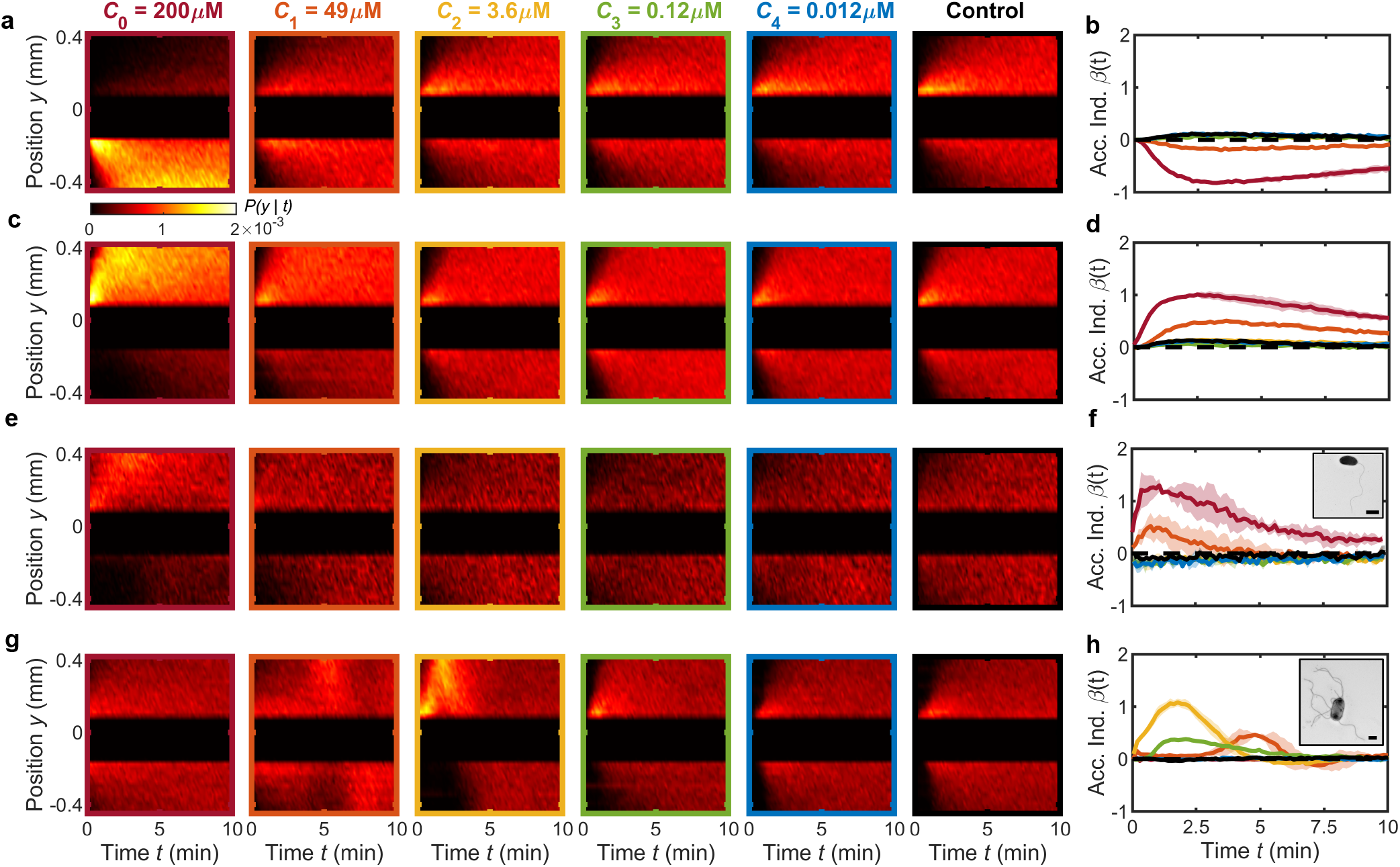
MCD enables rapid quantification of chemotactic responses across different chemostimulants and microbial species. (**a,b**), The negative chemotactic response of *V. alginolyticus* to the repellent phenol [37] is evident in kymographs of cell position, *P*(*y*|*t*), and the accumulation index, *β*(*t*), respectively. Central 250 *μ*m wide band of non-motile cells is omitted from *P*(*y*|*t*) for clarity with no impact on *β*(*t*). (**c,d**), MCD measurements demonstrate the positive chemotactic response of *V. alginolyticus* to leucine observed in *P*(*y*|*t*) and *β*(*t*), respectively. (**e-h**), *P*(*y*|*t*) in response to various concentrations of serine for bacteria *P. haloplanktis* (e,f) and *E. coli* (g,h). TEM images of *P. haloplanktis* (f, inset) and *E. coli* (h, inset). Scale bars, 1 *μ*m. *β*(*t*) for *P. haloplanktis* (f) illustrates a monotonically increasing response to increased concentrations (extracted from e). Accumulation index, *β*(*t*), for *E. coli* (h) reveals a peak response to intermediate serine concentration (*C*_2_ = 3.6 *μ*M) and delayed accumulation at higher concentrations [38]. Shaded regions are standard deviation (N=2 and N=3 for *P. haloplanktis* and *E. coli*, respectively). Color bar corresponds to kymographs in a,c,e,g.

To further illustrate the capabilities and flexibility of the MCD, the chemotactic behavior of *Pseudoalteromonas haloplanktis* (Fig. 3f, inset) and *Escherichia coli* (Fig. 3h, inset) to serine was measured (Fig. 3e,g) with no design changes to the MCD (see **Methods**). *P. haloplanktis* is a rapid-swimming, monotrichous marine bacterium that is a model organism for chemotaxis to amino acids [44] and exhibits strong chemotaxis towards cellular exudates [45]. Similar to *V. alginolyticus*, it utilizes a run-reverse-flick foraging strategy [46] for efficient chemotaxis in patchy chemical landscapes [6]. *P. haloplanktis* exhibited a monotonically increasing chemotactic response (Fig. 3f) with increasing serine concentration, qualitatively comparable to the response of *V. alginolyticus* to leucine (Fig. 3d). In contrast to the marine prokaryotes, *E. coli* is a pathogenic bacterium, which uses the bundling and unbundling of its multiple flagella to perform run-tumble motility for migrating up or down chemical gradients [1, 2]. *E. coli* has served as the canonical organism for bacterial motility and chemotaxis [1, 2, 18, 47] and has been instrumental in our understanding of logarthimic-sensing [10, 17] and chemical navigation in complex physical environments [38]. A single MCD assay reveals that *E. coli* has a strong chemotactic response to intermediate serine concentrations (*C*_2_; Fig. 3g,h). The response significantly diminishes for high (*C*_0-1_; Fig. 3g,h) and low chemostimulant concentrations (*C*_3-4_; Fig. 3g,h). This result is consistent with *E. coli*’s affinity for serine [39], but toxicity at high concentrations [48]. Furthermore, at higher concentrations (*C*_1_), the initial accumulation is delayed in time (Fig. 3h), an alluded feature of the chemotactic sensitivities of this organism to serine [38, 46]. Taken together, these three model swimming chemotactic microbes cover diverse foraging and motility strategies, whose range of chemotactic responses were efficiently screened using the MCD. These results demonstrate that the MCD can rapidly ascertain chemotactic responses across different chemostimulants and concentration ranges, which will be particularly valuable for studying the nano-molar and even femto-molar concentrations that characterize the detection thresholds of many microorganisms [11, 19, 32].

Our ability to simultaneously quantify a microbe’s response across a spectrum of attractant concentrations using the MCD now enables rapid comparative studies across microbial or chemical species (Fig. 4). The magnitude of a microbe’s response to a given concentration gradient is compactly summarized by the maximum (or minimum for negative chemotaxis) of their accumulation index, *β_max_* = ±max(|*β*(*t*)|), where the sign of *β_max_* is determined by the sign of the chemotaxis. In the case of serine, *V. alginolyticus* and *E. coli* (Fig. 4) both appear to have developed a chemotactic affinity to an optimal concentration [10, 38], which is evident from the local maxima of *β_max_* and is intrinsically linked to their motility and sensing abilities. Despite its many similarities to *V. alginolyticus, P. haloplanktis* (Fig. 4) exhibits a monotonically increasing response to serine across the concentrations tested. Separately, the lack of an optimal concentration in the response of *V. alginolyticus* to leucine likely indicates a higher-saturation threshold in the relevant chemoreceptors in comparison to its response to serine. This demonstrated screening efficiency highlights the benefits of this new microfluidic platform for tackling large-scale chemotaxis studies [20], regardless of the range of organisms or compounds of interest.

**FIG. 4.**
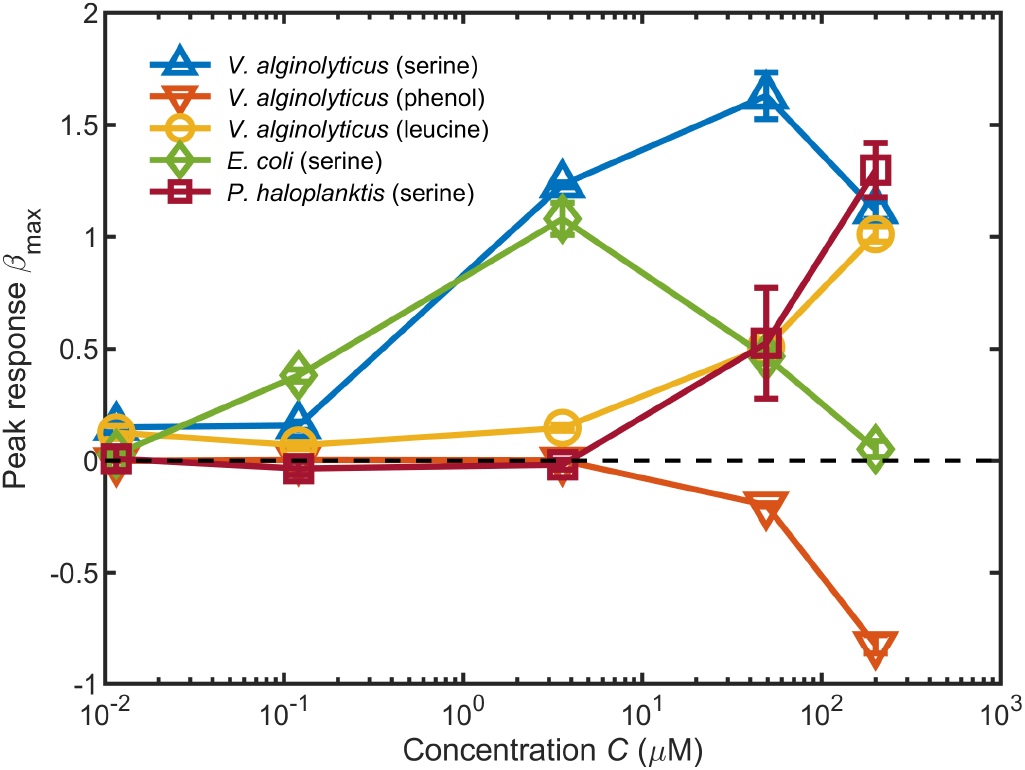
Summary of chemotactic responses measured using the MCD. Chemotactic responses were measured across various species, chemostimulants, and concentrations using the MCD. These responses are quantified by the peak of the accumulation index *β_max_* = ±max(|*β*(*t*)|), where the sign is determined by the positive or negative chemotactic behavior for each chemostimulus concentration (from Figs. 2h and 3b,d,f,h).

## III. DISCUSSION

Here, we have introduced a novel microfluidic multiplexed chemotaxis device for rapid quantification of bacterial responses to a range of chemostimulant concentrations. Identifying the diverse chemical compounds and concentrations responsible for driving microbial interactions that underpin important environmental and human health processes – for example, ecosystem scale nutrient cycling and disease transmission – has proven to be a tedious and monumental undertaking. A primary challenge is the sheer diversity of potential compounds and the extensive concentration range of microbial responses from micro- to femtomolar [11, 19, 32]. Unlike conventional microfluidic chemotaxis assays that are ill-equipped to address these challenges [20], our new multiplexed chemotaxis device (MCD) design mitigates cumbersome assays by rapidly screening the chemotactic behavior of microbes across a spectrum of chemostimulus conditions simultaneously (Fig. 1f). This work opens new avenues to large-scale panel experiments previously inaccessible with existing microfluidic devices.

The MCD’s two-layer device architecture uses a single pressure source to serially dilute a chemostimulus with a buffer input, producing five logarithmically separated chemical solutions (Fig. 1j). A cell suspension and additional motility buffer are introduced into each of six observation regions, along with the precisely controlled chemostimulus solutions, where the chemotactic response of the cells is recorded. The simultaneous chemotaxis assays are not only fast and efficient compared to conventional devices, but they also eliminate variability due to multiple culture preparations and aging cells (Fig. 2g). Taken together, our results illustrate that the MCD enables robust and efficient quantification of the chemotactic responses of various bacterial species to different chemostimulants, simplifying the labor-intensive chemotaxis screening process (Fig. 4).

The MCD design is amenable to a range of modifications to tailor its application, including but not limited to, alteration of chemical concentrations, gradients and flows, increased multiplexing, and cell retrieval. The sensitivity of prokaryotes is intrinsically linked to the strength of the concentration gradient [10, 11, 17, 47], an effect which can easily be examined with the MCD. With no alterations to the design or operation, simply changing the concentration of the input chemostimulus (*C*_0_, Fig. 1d) will shift the examined concentration range as desired. Likewise, the background concentration across all observation channels is easily adjustable by exchanging the buffer solution inputs for a non-zero concentration chemostimulus solution. The serial dilution layer hydraulic circuit design can be modified to produce specific dilution ratios (i.e. linear or logarithmic) [15, 31], where optimization of the device layout can expand the number of different chemical conditions probed. The MCD currently operates with a single outlet for operational simplicity, but sub-populations of chemotaxing cells could be captured for further analysis such as sequencing [11, 49]. The device can also be operated in a steady flow regime [13, 14] to study the growth and adhesion of cells in the presence of different chemical gradients in the observation regions, for example relevant to biofilm formation [50]. While the work presented here focuses solely on prokaryotes, the current device geometry will accommodate larger eukaryotic cells (≈ 10 *μ*m) and may be scaled up for multicellular microorganisms [51].

In summary, the MCD can accelerate discovery in both ecosystem and human health investigations. The engineered microfluidic device presented here will simplify the study of microbial chemotaxis, which is paramount to understanding and modeling global scale carbon and nutrient cycling [21] on a changing planet. Additionally, microfluidics has been identified as a potential means for meeting the high-throughput demands of chemical synthesis, screening, and testing with living cells, that remain key issues in drug discovery [52] to meet the challenges of antibiotic resistant microbes [53].

## METHODS

### A. Microfluidic device design

#### 1. Hydraulic circuit framework

In analogy with electrical circuits, well-established hydraulic circuit theory [54] was used to design the complex microfluidic network of the multiplexed chemotaxis device (MCD; see Fig. 1 and Fig. 6). Briefly, for incompressible, laminar flow through a constant cross-section microchannel, the pressure drop, Δ*p*, is linearly proportional to the volumetric flow rate, *Q*, and is given by Hagen-Poiseuille law [54, 55]: Δ*p* = *QR*. The hydraulic resistance, *R*, is a function of the fluid viscosity (properties of water assumed for all fluids) and the channel geometry. Fabrication of microfluidic devices via the soft lithography method [56] used here (see below) results in rectangular cross section microchannels (height, *H*; width, *W*; length, *L*). Exact expressions for R are tabulated for rectangular and other cross-section channels and provided in various resources [57]. Combined with conservation of mass, Σ*Q_i_* = 0 at the junctions (nodes) between several channels, *i*, the Hagen-Poiseuille law enables us to design complex microfluidic networks (Fig. 6) via the solution of a set of linear equations.

#### 2. MCD design considerations

The primary goal of the MCD was to efficiently perform several stop-flow bacterial chemotaxis assays [13, 58] simultaneously for a range of chemostimulus concentrations. The design requirements were to: (i) dilute and distribute five logarithmically spaced concentrations of chemostimulus plus one control buffer solution to each of six chemotaxis assays. (ii) Perform those six chemotaxis assays in parallel on the same microfluidic chip. And, (iii) the microfluidic device should receive minimal fluid inputs to reduce setup time. The MCD has a two-layer architecture (dilution layer and cell injection layer) with a total of four fluid inputs and one (waste) output (Fig. 1 and Fig. 6). Each having two inputs, the dilution layer and cell injection layer are designed to be regulated by a pressure-driven flow controller operating at a pressure, *p*_*in*,1-2_ ≈ 100mbar and *p*_*in*,3-4_ ≈ 50mbar, respectively, while the lone output is at atmospheric pressure (*p_out_* = 0). The dilution layer receives a base concentration chemostimulus solution (concentration, *C*_0_; *p*_*in*,1_) and a buffer solution (*C* = 0; *p*_*in*,2_), and the cell injection layer receives a bacterial suspension in buffer (*p*_*in*,3_) and a second buffer solution (*C* = 0; *p*_*in*,4_). The serial dilution [15] process sequentially combines the chemostimulus and buffer fluids to produce separate microchannel streams having chemostimulus concentrations of *C_i_* = 10^-*i*^*C*_0_ (*i* ∈ [0, 4]) and *C*_5_ =0 (control). The resulting six diluted chemostimulus solutions are merged in separate observation channels with a flow-stratified bacterial suspension and second buffer stream, which eventually forms the chemostimulus gradient in the chemotaxis assay. The three solutions are designed to symmetrically occupy the following fractional widths of the observation channel (total width, W): *w*_1_ = 4*W*/9, *w*_2_ = *W*/9, and *w*_3_ = 4*W*/9 (Fig. 6b). The observation channel width (*W* = 1 mm) and height (*H* = 90 *μ*m) were chosen to set the chemostimulus gradient strength based on a physically relevant length scale [13, 28, 47] and to ensure that the upper and lower microchannel walls do not impede bacterial motility, respectively. These dimensions are in line with conventional microfluidic chemotaxis assays [13], and as a consequence of the microfabrication process, the chosen H sets the height of the dilution layer channels excluding the micromixer. The initial flow rate of the three streams in the observation region (Fig. 1d) was designed to be 20, 5, 20 *μ*l min^-1^ for the chemostimulus (*Q_out_*), cell (*Q*_*CIL*,1_), and buffer solution (*Q*_*CIL*,2_), respectively (Fig. 6). This 4:1:4 flow rate ratio maintains stratification of cells, while ensuring that the bacterial cells are not damaged by the flow. Beyond these design requirements, the geometries - and thus hydraulic resistances – of several components are set independently, including: micromixer channels *R_M_*, bridge channels *R_B_*, and observation channels *R*_4,4_ (Fig. 6 and Table I). Based on these design requirements, hydraulic circuit theory was used to determine the required resistances of each microchannel in the MCD network, and subsequently the microchannel geometries [54]. A complete list of the microchannel resistances and dimensions for the final design is provided in Table I.

**TABLE I.** Multiplexed chemotaxis device (MCD) microchannel resistance and dimensions. Hydraulic circuit analysis (Methods) was used to determine the hydraulic resistance and thus geometry of the individual microchannels comprising the MCD channel network (Fig. 1d,e). The location of each channel in the dilution layer is indicated in the device layout in Fig. 1d and in the hydraulic circuit in Fig. 6a. Subscripts “CIL” indicate channels in the cell injection layer shown in Fig 1e and Fig. 6b. The resistance for the observation region (*R*_4,4_) varies in the microchannel width due to the hydraulic resistance varying between the channel sections before and after the chemical and buffer solution meet (Fig. 1d), and the micromixer (*R_M_*) has nonuniform channel height due to the ridges (**Methods**; see also Fig. 5)

| Channel | Resistance<br>(mPa·s·$\mu\text{m}^{-3}$) | Height<br>($\mu\text{m}$) | Width<br>($\mu\text{m}$) | Length<br>(mm) |
| --- | --- | --- | --- | --- |
| $R_{0,2}$ | 0.0082 | 90 | 90 | 19.06 |
| $R_{1,1}$ | 0.0095 | 90 | 90 | 22.08 |
| $R_{2,1}$ | 0.0147 | 90 | 90 | 34.06 |
| $R_{3,1}$ | 0.0201 | 90 | 90 | 46.43 |
| $R_{4,1}$ | 0.0277 | 90 | 90 | 64.13 |
| $R_{5,1}$ | 0.0300 | 90 | 90 | 69.45 |
| $R_{0,4}$ | 0.0201 | 90 | 90 | 46.50 |
| $R_{1,4}$ | 0.0149 | 90 | 90 | 34.54 |
| $R_{2,4}$ | 0.0098 | 90 | 90 | 22.60 |
| $R_{3,4}$ | 0.0046 | 90 | 90 | 10.76 |
| $R_B$ | 0.0035 | 90 | 150 | 19.93 |
| $R_{\text{CIL},1}$ | 0.0575 | 75 | 75 | 64.24 |
| $R_{\text{CIL},2}$ | 0.0142 | 75 | 75 | 15.89 |
| $R_M$ | 0.0043 | - | 200 | 40.7 |
| $R_{4,4}$ | $7.7 \times 10^{-4}$ | 90 | - | - |

### B. Herringbone micromixer design

#### 1. Mixing performance

For the serial dilution process to perform as designed, effective mixing of the chemical solution and buffer are critical. Here, we use a well-established herringbone micromixer geometry [29], where a series of ridges on the upper wall of an otherwise rectangular microchannel (Fig. 5a,b) drive a three-dimensional flow to enhance mixing [29, 59]. A separate microchannel - having the same cross section geometry as the MCD design - was fabricated to independently quantify mixing performance and to select the necessary mixer length. The test micromixer channel was 41.3 mm long with 29 mixing cycles (comprised of two sets of six alternating herringbone ridges each). Two aqueous solutions of fluorescein salt (Sigma; concentrations, *C*_50_ = 50 *μ*M and *C*_10_ = 10 *μ*M) [60] were injected individually into the MCD, and calibration images of dye intensity were captured after each herringbone mixer cycle (Fig. 5), corresponding to the maximum (**I**_50_) and minimum (**I**_10_) dye concentrations, respectively. The region within 20 *μ*m of the microchannel walls was excluded from analysis due to reflection and refraction effects [29]. Subsequently, the two solutions were flowed side-by-side with images (**I**_*i*_) recorded in the same locations as above and normalized as follows:

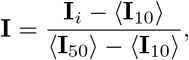

where 〈·〉 denote spatial averaging. The degree of mixing is defined as [29], 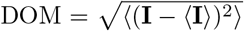, where values of 0.5 and 0 indicate fully non-mixed and mixed solutions, respectively. This measurement (Fig. 5c) was repeated for both the designed flow rate for the MCD (*Q_M_* = 22 *μ*l min^-1^) and for a second higher flow rate (222 *μ*l min^-1^). Based on standard metrics [29], the two solutions are considered mixed when DOM ≤ 0.05 (i.e. 90% complete mixing). For both flow rates, this criterion is met after 9 complete herringbone ridge cycles, and a final design with 26 herringbone cycles was chosen for the MCD. The independence of mixing efficiency on flow rate, combined with a safety factor of approximately three for the number of herringbone cycles, ensures that the serial dilution portion of the MCD will perform accurately for a wide range of chemostimulants and flow speeds.

**FIG. 5.**
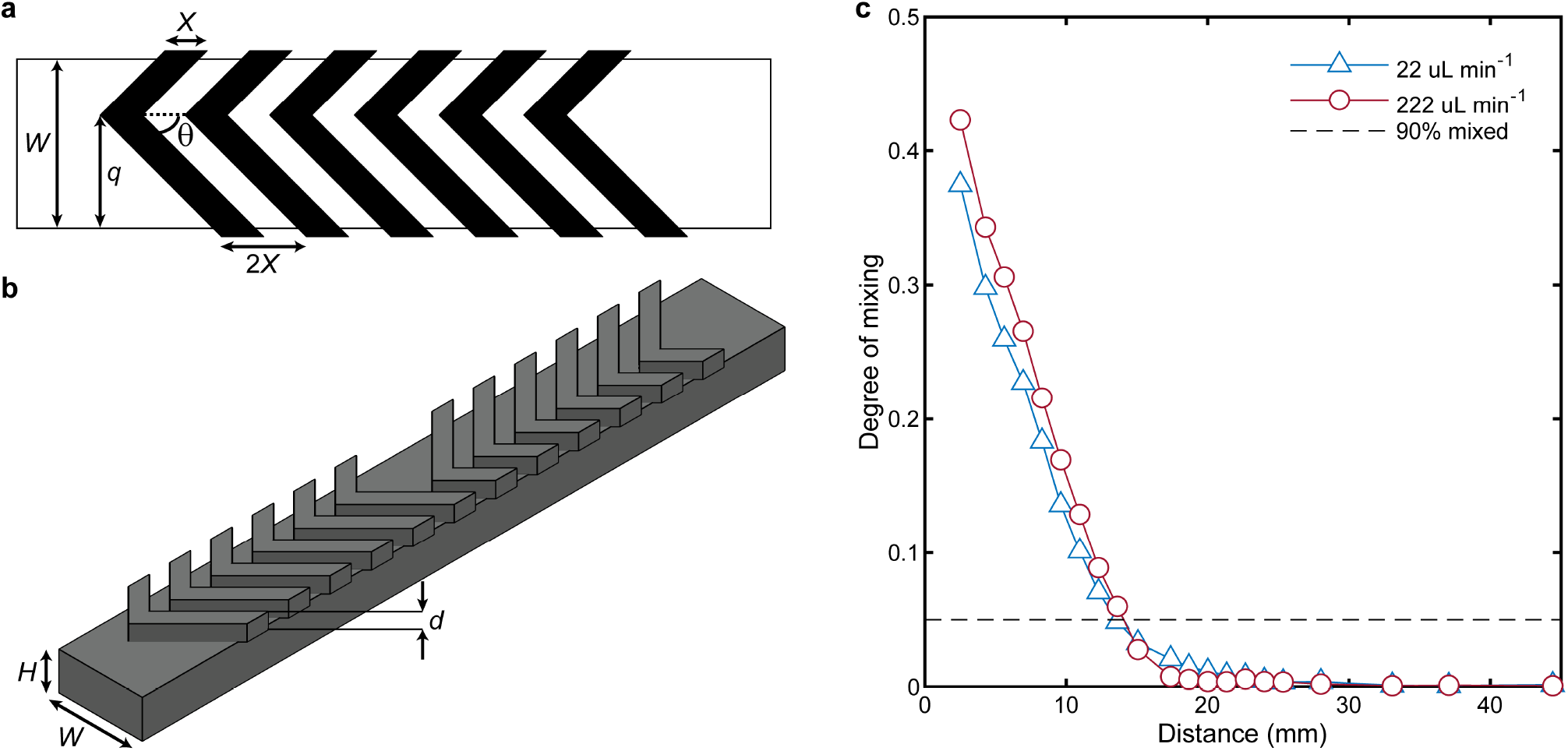
Micromixer geometry and mixing performance. (**a,b**), The herringbone micromixer [29] - used to incorporate chemical and buffer during serial dilution - consists of a main rectangular channel (*W* = 200 *μ*m, *H* = 90 *μ*m) with herringbone ridges (*X* = 50 *μ*m, *d* = 30 *μ*m), which enhance mixing by generating transverse flow. Each herringbone half cycle (≈ 700 *μ*m long) consists of six ridges (pitch, *θ* = 45°; a). The distance, q, from the side wall alternates every half cycle between 2W/3 and W/3. (**c**), The degree of mixing [29] (DOM; Methods) was quantified for a long test channel having 29 herringbone cycles for two different flow rates: 22 *μ*l min^-1^ (blue triangles) and 222 *μ*l min^-1^ (red circles) using fluorescein dye [60] (*D* = 436 *μ*m^2^ s^-1^). Distance indicates the downstream position from the point where the dye and water solutions meet. The solution is considered mixed when DOM ≤ 0.05 (90% complete mixing, dashed line), which is achieved after 9 cycles for both flow rates. The final micromixer design comprised 26 herringbone cycles (see Figs. 6,8) to ensure complete mixing across a range of potential chemostimulants.

#### 2. Micromixer hydraulic resistance

To complete the design of the MCD, it was necessary to determine the hydraulic resistance of the herringbone micromixer *R_M_* which was measured empirically using a parallel microfluidic device [61]. Briefly, a microfluidic device was fabricated with two parallel channels connected by shared inlets and outlets. The parallel channels had identical rectangular geometries except one had the herringbone ridges replicating the micromixer channel section *R_M_* (Fig. 8). Two solutions, DI water and 1 *μ*l ml^-1^ tracer particle suspension (0.25 *μ*m radius; 2% solid; carboxylated FluoroSpheres, Life Technologies), were flowed through the device using glass syringes (2.5 ml; Hamilton) mounted on two separate syringe pumps (Harvard Apparatus). The particle solution was visualized using fluorescence microscopy, and the flow rates of the two pumps were adjusted such that the two streams divided equally into the parallel channels. The micromixer hydraulic resistance was determined from the resulting flow rate ratio and the known (analytical) resistance of the non-mixer channel [54] (*R_M_* = 0.0043 mPas *μ*m^-3^; Fig. 6,8c), and the results were corroborated by COMSOL Multiphysics simulations (not shown).

### C. Microfabrication

Microfluidic channel molds were fabricated using standard single and two-layer photolithography [62] to transfer the final channel designs from a photomask (Artnet Pro, formally CAD/Art Services, Inc.) onto a silicon wafer (100 mm diameter; University Wafer), which was spin-coated with photoresist (SU-8; Kayaku Advanced Materials). The single assay chemotaxis devices and MCD cell injection layer were made using SU-8 2050 and 2025, respectively, and multilayer devices (micromixer validation channels, MCD dilution layer) were made using SU8-3050 and SU8-2025 for the main rectangular channels and herringbone ridges, respectively. The ridges of the micromixers [29] were applied by halting the first-layer photolithography after the first post-exposure bake (PEB), spin-coating a second layer of SU-8 photoresist onto the wafer, then completing the remainder of the photolithography processes as usual [62]. The ridges of the herringbone micromixer [29] extend over the main channel by ≈ 10 *μ*m on both sides to account for misalignment during the multilayer photolithography (Fig. 5, Fig. 8). As fabricated, the final channel heights for the MCD dilution layer (Fig. 1d and Fig. 8a-c) were 90 – 94.5 *μ*m, 37–38.5 *μ*m for the main channel and herringbone ridges, respectively, while the cell injection layer (Fig. 1e) was 71 – 73 *μ*m high (Bruker’s DekTak).

The MCD was fabricated using two-layer soft lithography [56] with polydimethylsiloxane (PDMS; Sylgard 184) at a 10:1 (elastomer:curing agent) ratio. All channel wells were punched using a 1.5 mm diameter biopsy punch (Integra). The dilution layer mold was first silanized through vapor deposition [63] in a vacuum desiccator with 1-2 drops of tridecafluoro-1,1,2,2-tetrahydrooctyl trichlorosilane (Gelest Inc.) to help release the cast PDMS. Post-silanization, PDMS was poured onto both the cell injection layer and dilution layer molds and degassed in a vacuum chamber (≈ 1 hour) prior to curing (65°C for ≈ 1 hour). The resulting PDMS dilution layer channel was first plasma bonded onto a standard thickness, double wide glass slide (75 mm × 50 mm × 1 mm; Fisherbrand) using a plasma oven (Plasma Etch Inc.), and subsequently heated on a hot plate at 110° C for one hour to promote covalent bonding [56] (Fig. 8d). Next, the cell injection layer was plasma bonded on top of the dilution layer, with care taken to ensure the alignment of the fluid wells connecting the two layers (Fig. 8e). The assembled device was baked again on a hot plate at 110° C for one hour. The PDMS-PDMS bond was found to be sufficiently strong for the relatively low pressure applications of the MCD [64]. All other devices (e.g., single assay chemotaxis devices and micromixer validation channels) were fabricated using single-layer soft lithography, where an individual PDMS device was molded and subsequently bonded to a standard microscope slide using the procedures described above.

### D. MCD dilution, flow, and gradient generation performance

The performance of the fabricated MCD was validated using epifluorescence microscopy (Nikon Ti-E) with an aqueous fluorescent dye (fluorescein sodium salt, Sigma) of various concentrations (described below), and images of the dye distribution were captured at the midplane of the channels with a sCMOS camera (Zyla 5.5; Andor Technology). Fluorescein was chosen due to its similar diffusion coefficient with the chemostimulant serine [32].

#### 1. Serial dilution

The primary function of the MCD dilution layer is to sequentially dilute the input chemical solution (*C*_0_) with buffer to generate four logarithmically decreasing concentrations (*C*_1-4_) for each of the observation channels (plus one control, *C*_5_ = 0). The dilution performance was quantified by injecting a solution with known fluorescein concentration (*C*_0_ = 1 mM). The diluted concentration field, *C_i_*(*x, y*), at each of the observation channels was then determined from the local measured dye intensity, **I**_*i*_(*x,y*), which are linearly proportional, *C_i_*(*x,y*) ∝ **I**_*i*_(*x,y*). Fluorescence images were recorded upstream of the inlet before the three fluids made contact in each observation channel (Fig. 1a) using a 20× (0.45 NA) objective. Due to the strong ten-fold dilution, pairs of images were acquired for adjacent channels with optimized exposure times to account for the finite dynamic range of the camera. The mean normalized concentrations provided for each observation region from the serial dilution process were reconstituted from the measured image intensity as follows:

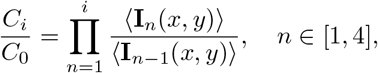

where the angled brackets indicate the spatial average. The resulting serial dilution followed the expected logarithmic (10-fold) dilution *C_i_*/*C*_0_ = 10^-*i*^ for *i* ∈ [0, 4] for which the system was designed (Fig. 1j). This measurement was performed for three different sets of applied driving pressures, which yielded nearly identical results and illustrated the robustness of the serial dilution process.

#### 2. Stratification symmetry

The symmetry of the stratified chemostimulus and buffer distributions in the observation channel is critical to prevent bias in the chemotaxis measurements. As minor errors in the manufacturing process can alter this symmetry, the widths of the chemical, cell, and buffer streams in each observation channel were measured by flowing a fluorescein solution (*C*_0_ = 100 *μ*M) in both the chemical and buffer inlets of the dilution layer as well as the buffer inlet of the cell injection layer (Fig. 1d,e). By flowing DI water only through the cell suspension inlet of the injection layer, a strong intensity contrast between the central cell band and the surrounding fluids was visualized in the observation channel (Fig. 7a). Images were recorded (10×, 0.3 NA objective), and the widths of the three fluid streams, *w_i_*, were measured based on their spatial intensity distribution (Fig. 7b). The normalized widths of the chemostimulus and buffer streams, 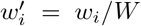, across all observation channels deviated from the original design width by only 2.6% and 1.8% on average, respectively, while the mean deviation in cell suspension stream width was just 3.0%.

**FIG. 6.**
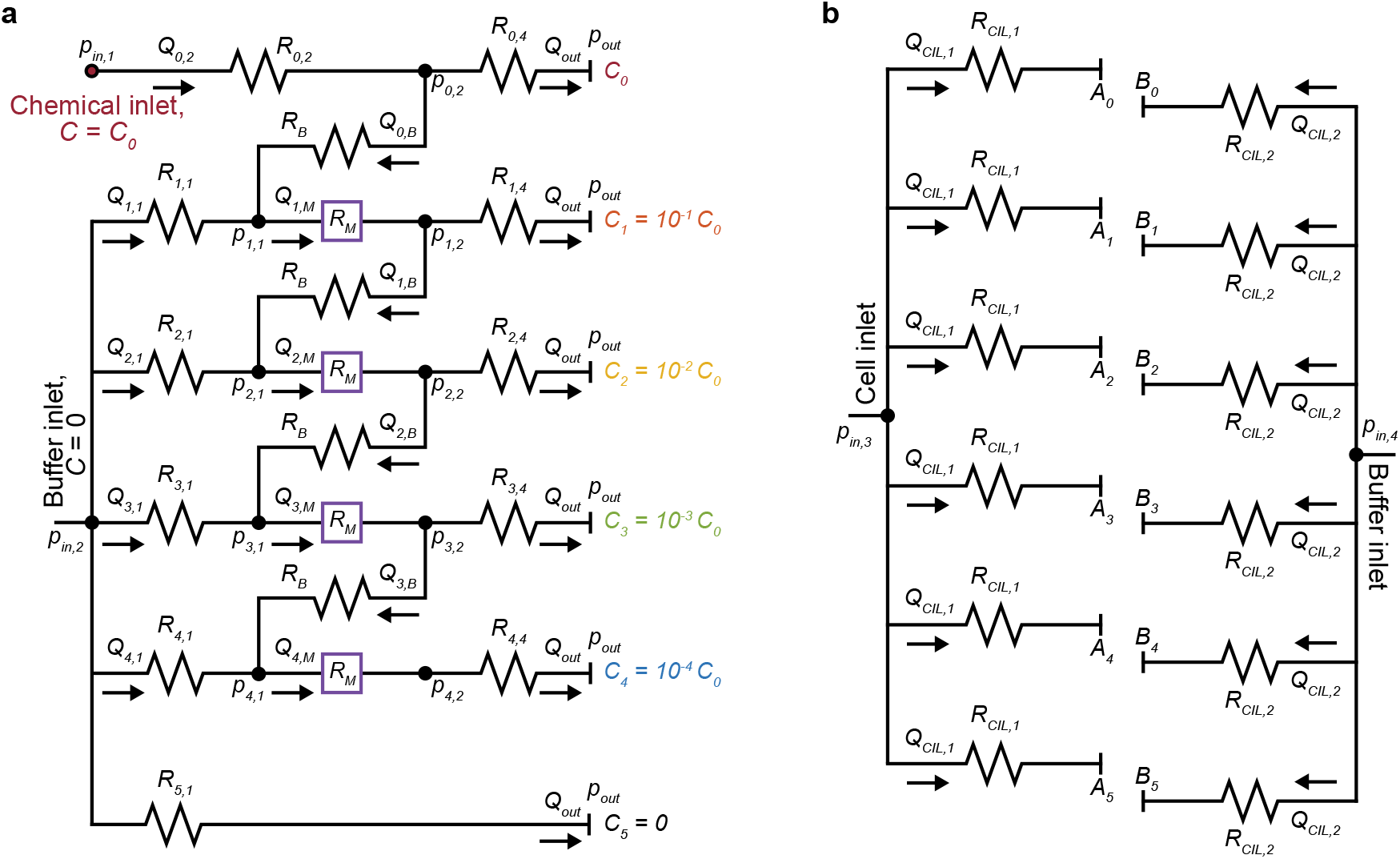
Hydraulic circuit design of MCD dilution layer and cell injection layer. (**a**), Circuit representation of the dilution layer [15] of the MCD (Fig. 1d). The two inlets (*p*_*in*,1_, *p*_*in*,2_) receive the base chemical solution (*C*_0_) and the buffer solution (*C* = 0), respectively, and the outlet is open to atmospheric pressure (*p_out_* = 0mbar). The hydraulic network was designed such that a prescribed inlet pressure *p*_*in*,1_ = *p*_*in*,2_ = 100mbar from a pressure controller produces outlet flow rates of *Q_out_* = 20 *μ*l min^-1^. The resistances of the bridge channels (*R_B_*), micromixer channels (*R_M_*), and observation channels (*R*_4,4_) were predetermined (Methods). The remaining flow rates, pressures, and hydraulic resistances were determined by solving a linear set of equations for the hydraulic circuit [55], and the resulting resistances determine the individual channel geometries (Table I). (**b**), Circuit representation of the cell injection layer of the MCD (Fig. 1e). The cell injection layer network was designed to flow the cell (*p*_*in*,3_) and buffer (*p*_*in*,4_) solution into the dilution layer to ensure the cells would be placed in the center of the observation region. To do so, the prescribed inlet pressures were *p*_*in*,3_ = *p*_*in*,4_ = 50mbar and flow rates in each microchannel for the cell (*Q*_*CIL*,1_) and buffer (*Q*_*CIL*,2_) were 5 *μ*l min^-1^ and 20 *μ*l min^-1^, respectively. The resulting channel resistances and dimensions were determined using Hagen-Poiseuille law [54, 55] (Methods), and accounted for the resistance of the observation region (*R*_4,4_; Table I). The labels in a and b corresponds to that in Fig. 1d,e.

**FIG. 7.**
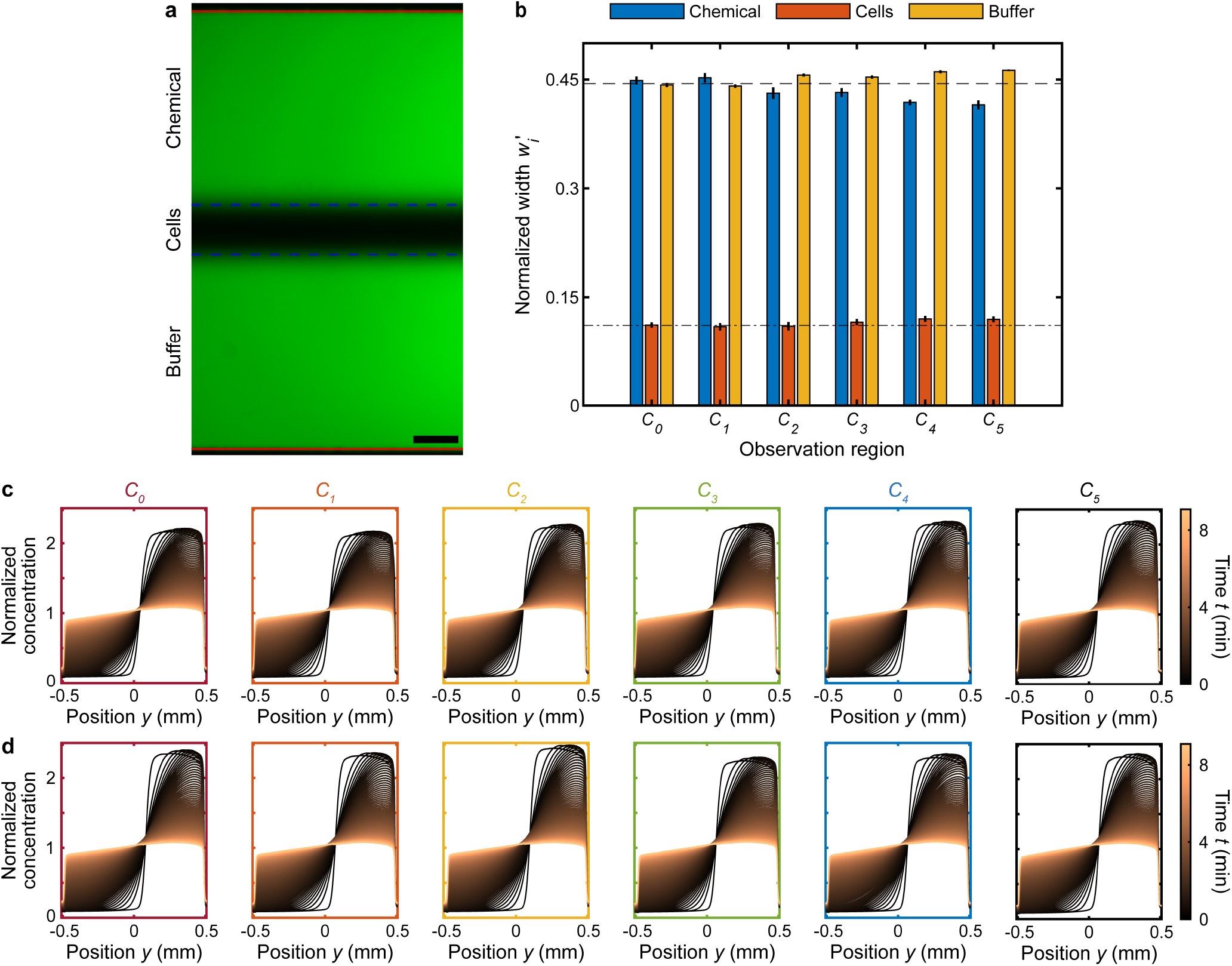
Validation of chemostimulus distribution and gradient evolution in MCD observation regions. (**a**), The initial symmetry of the stratified chemical, cell, and buffer solutions in the observation channel was verified by measuring the widths of the three fluid streams through fluorescent dye visualization (Methods). A fluorescein solution was flowed through both the chemical and buffer inlets of the dilution layer and also the cell solution inlet of the cell injection layer. Solid red and dashed blue lines indicate measured channel wall and stream interface locations, respectively, for a driving pressure *p*_*in*,1_ = *p*_*in*,2_ = 200 mbar and *p*_*in*,3_ = *p*_*in*,4_ = 140 mbar (Fig. 1d,e and Fig. 6). Scale bar, 100 *μ*m. (**b**), The normalized measured widths, 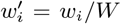, of the chemical, cell, and buffer streams illustrate their high degree of symmetry across all observation channels (*C*_0-5_); where *w_i_* is the width of fluid stream (a; dashed blue lines) and W the total width of the intensity profile (a; solid red lines). The bar plots and error bars indicate the mean and standard deviation of the normalized stream widths, investigated for three different driving pressures of the dilution layer and cell injection layer: 100/70 mbar, 150/105 mbar, and 200/140 mbar (*p*_*in*,1-2_/*p*_*in*,3-4_). The chemical and buffer solutions percent difference from the original design width by only 2.6% and 1.8% on average, respectively, while the mean percent difference in cell suspension stream width was just 3.0%. To achieve symmetric flow, the applied pressure was tuned. This variation in the designed (Fig. 6) and applied inlet pressures (*p_in,i_*, *i* ∈ [1, 4]) could be due to the nonuniform fabricated channel height (Methods) and variations in the inlet tubing lengths that supplied the solutions to the MCD. (**c,d**), The uniformity of the chemostimulus gradient evolution across the various observation regions (*C*_0-5_) was determined through fluorescent dye (fluorescein) visualization of stop-flow experiments (Methods). The same dye concentration was injected into both the chemical and buffer inlets of the dilution layer to produce the same base concentration (*C_i_* = *C*_0_) and therefore gradient in each observation region for ease of comparison. The fluorescence intensity acts as a proxy to quantify the concentration profile across each channel over the course of approximately 9 min (post flow, t = 0). Measurements were performed for two different driving pressures to ensure that the resulting gradients were insensitive to flow rate: 100/70 mbar (c) and 200/140 mbar (d).

**FIG. 8.**
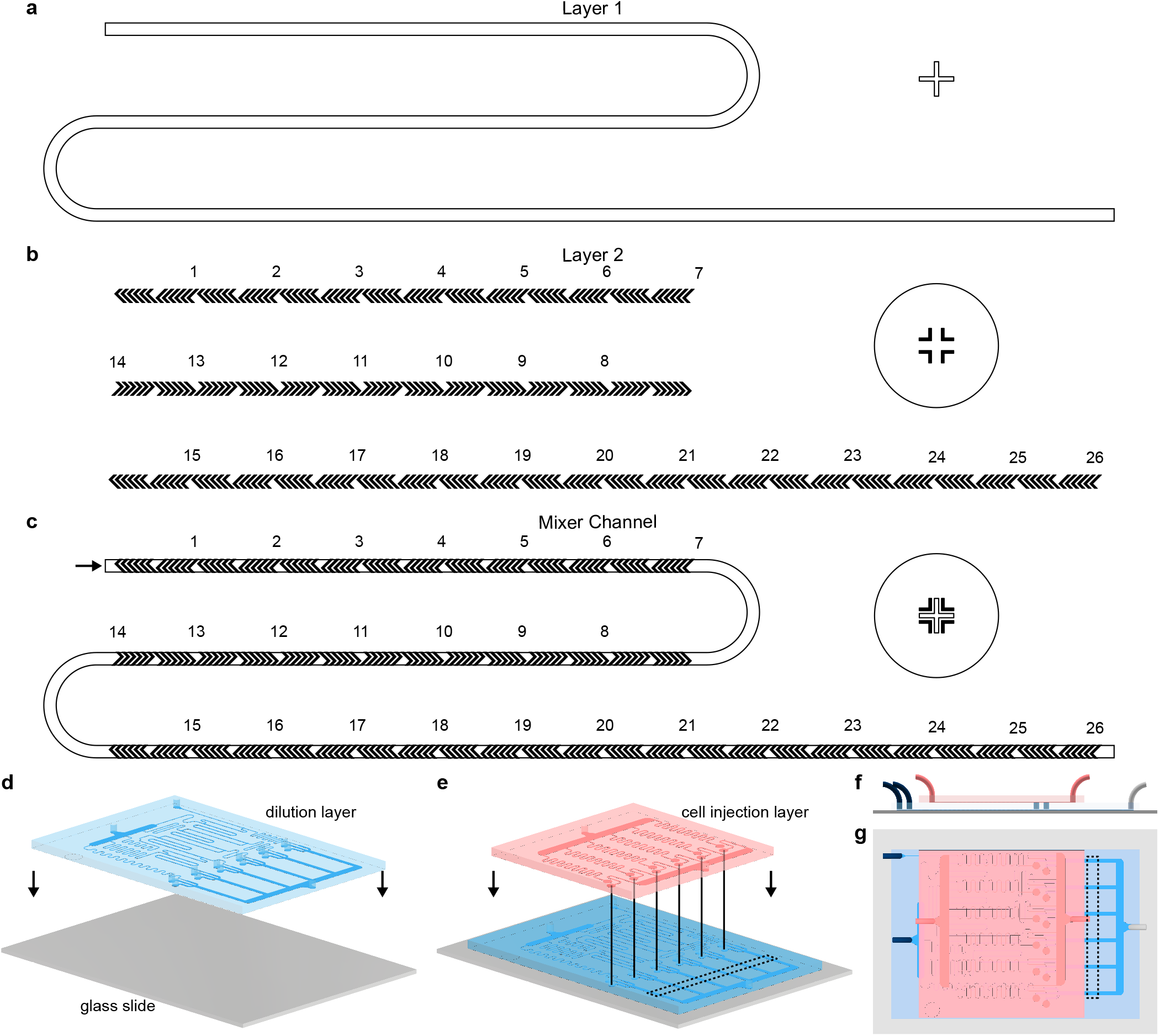
Two-layer photolithography and soft lithography microfabrication of the MCD. (**a-c**), Herringbone micromixers [29] were fabricated in the dilution layer of the MCD using multilayer photolithography (subsection of full photomasks shown; see also Fig. 1). The first photolithography layer produced the rectangular cross-section main channel (200 *μ*m wide; a), and a second photolithography layer formed the herringbone ridges on top [62] (b; see also Fig. 5). Complementary alignment markers were used to align the herringbone ridges (b, hollow cross and circle) with the main channel (a, cross). (**c**), Overlay of the herringbone micromixer channel components from a and b (see also Fig. 5). (**d,e**), The dilution layer (d, blue) and cell injection layer (e, red) PDMS microchannels for the MCD were cast individually from SU-8 molds via soft lithography [56] (Methods). The dilution layer microchannel was first plasma bonded [56] to a standard double-wide glass slide (d; grey; 75 mm × 50 mm × 1 mm). Subsequently, the cell injection layer was aligned and plasma bonded [56, 64] on top of the dilution layer (e). (**f,g**), Side and top-down view of the assembled MCD, respectively, showing the locations of the four inlets, single outlet, and observation regions (dashed box; see also Fig. 1).

**FIG. 9.**
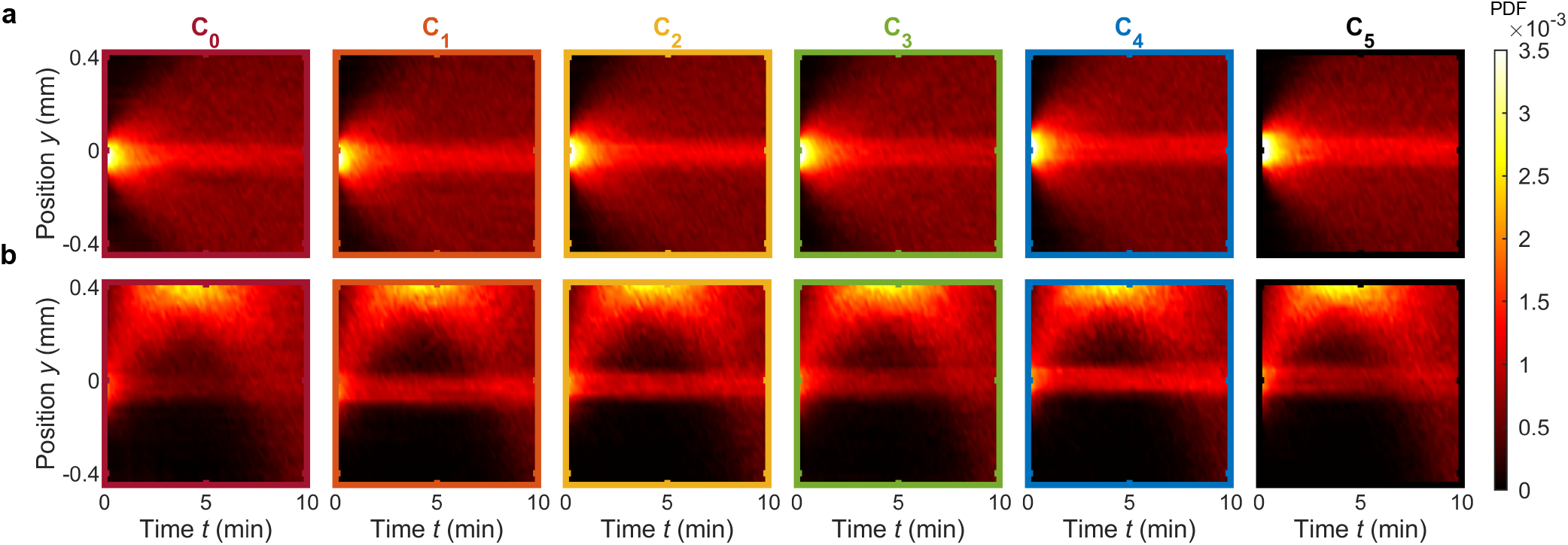
Control chemotaxis assays for different fixed gradient conditions. (**a**), Control chemotaxis assay with *V. alginolyticus* in the absence of a chemical gradient. In place of a chemostimulus, ASW (*C*_0_ = 0) was injected into the chemical inlet of the dilution layer, illustrating that bacteria exhibit no significant bias (see also Fig. 2k). (**b**), Control experiment with *V. alginolyticus* to a fixed chemical gradient across all observation channels illustrating consistent (positive) chemotaxis. ASW with serine at a concentration *C*_0_ = 200 *μ*M was injected into both the chemical and buffer inlets of the dilution layer. This approach results in each of the observation regions exhibiting the same chemostimulus gradient, based on the serine concentration *C_i_* = 200 *μ*M (see also Fig. 2l).

#### 3. Chemostimulus gradient consistency

Beyond ensuring the symmetry of the chemostimulus and buffer stratification, the time evolution of the resultant chemostimulus gradient must be consistent across each of the observation channels to accurately compare bacterial chemotactic responses. The chemical gradient evolution (Fig. 1k and Fig. 7c,d) was measured by first flowing a fluorescein solution (*C*_0_ = 100 *μ*M) through both the chemical and buffer inlets of the dilution layer and DI water through both inlets of the cell injection layer. Having independently verified the performance of the serial dilution process, this approach produces identical base concentrations for all observation channels, *C_i_* = *C*_0_, and thus, enables easy comparison of the resulting concentration gradients in each channel. Upon halting the flow, an image was recorded (10×, 0.3 NA objective) in each observation channel every 5 s for approximately 9 min. The time evolution of the (normalized) spatial fluorescence intensity was measured to visualize the chemical gradient. The resulting concentration profiles were found to be highly consistent across the various observation regions and for various driving pressures (Fig. 7c,d).

### E. Cell culturing

*Vibrio alginolyticus* (YM4; wild-type) from –80°C stock were grown overnight in Marine 2216 media (Difco) by incubating at 30°C and shaking at 600 revolutions per minute (RPM). The overnight culture was diluted 100-fold into fresh pre-warmed 2216 media and grown for three hours (30°C, shaking at 600 RPM) to O.D. ≈ 0.2. 7 ml of culture was then washed and resuspended (1,500 RCF for 5 min) in 4 ml of artificial seawater (ASW).

*Psuedoalteromonas haloplanktis* (ATCC 700530) from –80°C stock were grown overnight in Marine 2216 (Difco) media by incubating at room temperature and shaking at 100 RPM [6].

*Escherichia coli*(MG1655) from –80°C LB stock were grown overnight in Terrific Broth (TB, Sigma Aldrich) by incubating at 34° C and shaking at 220 RPM [6]. The overnight culture was diluted 100-fold into fresh pre-warmed TB media, and grown for approximately three hours (34°C, shaking at 220 RPM) to O.D. ≈ 0.5. 8 ml of culture was then washed three times and resuspended (4,000 RCF for 5 min) in 4 ml of motility buffer (10 mM potassium phosphate, 0.1 mM EDTA, 10 mM NaCl, pH 7, filter sterilized 0.2 *μ*m.). 16 ml of culture was washed twice and resuspended (1,200 RCF for 5 min) in 6 ml of ASW.

### F. Media and chemostimulants

Artificial seawater (ASW) was prepared following the NCMA ESAW Medium recipe, which was adapted from [65] and modified [66]. ASW was used as the buffer and the chemical solvent for chemotaxis assays for both *V. alginolyticus* and *P. haloplanktis*, while motility buffer (see above) was used for chemotaxis assays with *E. coli*. Chemostimulus materials were purchased from Sigma Aldrich for use in the chemotaxis experiments, specifically: serine (S4500), phenol (P1037), and leucine (L7875).

### G. Microfluidic chemotaxis assays

Prior to use, the MCD was pre-treated by flowing a 0.5% (w/v) bovine serum albumen solution (BSA; Sigma Aldrich) to reduce cell adhesion to the microchannel surfaces. The device was flushed for over 10 min prior to first use with the cell, chemostimulus, and buffer suspensions. For chemotaxis assays, fluid flow was driven by a single pressure controller (Elveflow OB1; 1 mbar = 100Pa): *p*_*in*,1-2_ = 200mbar (dilution layer) and *p*_*in*,3-4_ = 140mbar (cell injection layer). Pressures were scaled down to 100 mbar and 70 mbar, respectively, for *P. haloplanktis* experiments. Between each chemotaxis assay, the fluid inputs were flowed for a minimum of 2 min to stratify cell, chemostimulus, and buffer streams in the observation channels. Upon halting the flow, a monotonic concentration profile was established in each observation channel due to the diffusion of the chemostimulus (Fig. 1 and Fig. 7). The spatio-temporal evolution of the bacterial distribution was determined by imaging the cells using phase-contrast microscopy (10×, 0.3 NA objective; Tikon Ti-E) with a sCMOS camera (Zyla 5.5, Andor Technology) for approximately 10 min. An automated computer-controlled stage was used to cyclically move the microscope field of view to each observation channel 75 times, producing an effective imaging period of 8 s for each observation channel. Each experiment was replicated at least three times with the same culture and repeated on different days with freshly grown cells. For validation of the MCD, a conventional single assay (SA) microfluidic device (Fig. 2b-f) was designed with a similar geometry to the individual MCD observation channels. Specifically, the SA devices had three inlets (width, 0.5 mm) which merged in a single observation channel. The devices were fabricated in a single layer using soft lithography (see above), and they were pre-treated with a 0.5% (w/v) BSA solution and flushed with ASW prior to experiments. The SA chemotaxis devices were used to validate the MCD for the well-established chemotactic behavior of *V. alginolyticus* to the chemoattractant serine (Sigma). The three inlets of the single assay device (Fig. 2b) carried the chemoattractant dissolved in ASW, *V. alginolyticus* suspended in ASW, and ASW, respectively. The three solutions were flow stratified for a minimum of 2 min using a syringe pump (Harvard Apparatus), whereby flow rates were controlled by syringe size to maintain a 4:1:4 ratio of the stream widths. In an identical manner to the MCD, a chemostimulus gradient develops in the channel via diffusion, and the chemotactic response of the cell population was observed over time. Imaging was performed with phase-contrast microscopy (4×, 0.13 NA objective; Nikon Ti-E) at 1 fps over the course of approximately 10 min using a CMOS camera (Blackfly S, Teledyne FLIR).

### H. TEM imaging

For each species, initial cultures were grown following the previously described protocols (without any initial washing/resuspending), before the following final cell suspensions were prepared: (i) 4 ml of *V. alginolyticus* culture washed and resuspended (1,500 relative centrifugal force (RCF) for 5 min) in 1 ml of fresh 2216 media, (ii) 1 ml of *P. haloplanktis* culture washed and resuspended (1,200 RCF for 5 min) in 1 ml of fresh 2216 media, then diluted 10× in DDW (double distilled water), and (iii) 8 ml of *E. coli* culture washed and resuspended (4,000 RCF for 5 min) in 1 ml of DI water, then diluted 10× in DDW. For each species, 4 *μ*l of cell suspension was applied to a glow discharged copper mesh carbon coated grid and allowed to adsorb to the grid for 30 s. The grids were briefly washed in DDW, followed by staining with 1% Aqueous Uranyl Acetate, and allowed to dry fully before imaging. Grids were imaged using a FEI Morgagni transmission electron microscope (FEI, Hillsboro, OR) operating at 80 kV and equipped with a CMOS camera (Nanosprint5, AMT).

## AUTHOR CONTRIBUTIONS

M.R.S, R.J.H., and J.S.G. designed the research and the device; M.R.S fabricated the devices, performed validation assays, and analyzed corresponding data; R.J.H. performed chemotaxis assays and analyzed corresponding data; M.R.S, R.J.H., S.A.F., and J.S.G. discussed the results and wrote the paper.

## ACKNOWLEDGMENTS

We thank J. Vlahakis of the Tufts Micro and Nano Fabrication Facility for assistance in the device fabrication, J.B. Noble for preliminary work on the microfluidic device design, and R. Stocker and M. Salek for helpful discussions. TEM samples were prepared and imaged by the Brandeis Electron Microscope Facility. This work was funded by NSF Awards OCE-1829827, CAREER-1554095, and CBET-1701392 (to J.S.G), and OCE-1829905 (to S.A.F.).

